# TurboID mapping reveals exportome of secreted intrinsically disordered proteins in the transforming parasite *Theileria annulata*

**DOI:** 10.1101/2023.12.14.571664

**Authors:** Francis Brühlmann, Carmen Perry, Charlotte Griessen, Kapila Gunasekera, Jean-Louis Reymond, Arunasalam Naguleswaran, Sven Rottenberg, Kerry Woods, Philipp Olias

## Abstract

*Theileria annulata* is a tick-transmitted apicomplexan parasite that gained the unique ability among parasitic eukaryotes to transform its host cell, inducing a fatal cancer-like disease in cattle. Understanding the mechanistic interplay driving this transformation between the host cell and malignant *Theileria* species requires the identification of responsible parasite effector proteins. In this study, we used TurboID-based proximity labelling, which unbiasedly identified secreted effector proteins within host cell compartments. By fusing TurboID to nuclear export or localization signals, we biotinylated proteins in the vicinity of the ligase enzyme in the nucleus or cytoplasm of infected macrophages, followed by mass spectrometry analysis. Our approach revealed with high confidence nine nuclear and four cytosolic candidate effector proteins within the host cell compartments, eight of which had no orthologues in non-transforming *T. orientalis*. Strikingly, all eight of these proteins are predicted to be highly intrinsically disordered proteins (IDPs). We discovered a novel tandem arrayed protein family, Nuclear Intrinsically Disordered Proteins (NIDP) 1 - 4, featuring diverse functions predicted by conserved protein domains. Particularly, NIDP2 exhibited a biphasic host cell-cycle dependent localization, interacting with the EB1/CD2AP/CLASP1 parasite membrane complex during mitosis and the tumor suppressor Stromal Antigen 2 (STAG2), a cohesion complex subunit, in the host nucleus. In addition to STAG2, numerous NIDP2-associated host nuclear proteins implicated in various cancers were identified, shedding light on the potential role of the *T. annulata* exported protein family NIDP in host cell transformation and cancer-related pathways.

**IMPORTANCE:** TurboID proximity labelling was used to unveil the secreted proteins of *Theileria annulata*, an apicomplexan parasite responsible for a fatal, proliferative disorder in cattle, representing a significant socio-economic burden particularly in north Africa, central Asia, and India. Our investigation has provided important insights into the unique host-parasite interaction, revealing effector proteins characterized by high intrinsically disordered protein (IDP) structures. Remarkably, these proteins are conspicuously absent in non-transforming *Theileria* species, strongly suggesting their central role in the transformative processes within host cells. In addition, our study identified a novel tandem arrayed protein family, with Nuclear Intrinsically Disordered Protein (NIDP) 2 emerging as a central player interacting with established tumor genes. Significantly, this work represents the first unbiased screening for exported effector proteins in *Theileria* and contributes essential insights into the molecular intricacies behind the malignant transformation of immune cells.

## INTRODUCTION

The Apicomplexan phylum harbors diverse parasites, among which *Plasmodium*, *Toxoplasma*, *Cryptosporidium*, and *Theileria* stand out for their impact on human and animal health. Apicomplexans use sophisticated mechanisms to manipulate host cells, inducing metabolic shifts and changes in host gene expression (1). Transforming *Theileria* species orchestrate profound host cell changes that resemble cancerous cell phenotypes (2–4). The resulting disease in cattle, prevalent in the Southern Hemisphere and parts of Asia, is characterized by fever, anemia, and massive lymph node enlargements mirroring lymphoma, with a high mortality rate and a devastating impact on local farming communities (5–8). Despite similarities to cancer, including uncontrolled proliferation, invasiveness, and metastasis, the host cell DNA remains unaltered (9), making the intricate parasitic manipulation of basic cell biological signaling pathways in the host an intriguing system to study. In this study, we focused on *T. annulata*, the causative agent of Tropical Theileriosis in cattle, which infects macrophages and B cells that undergo extensive post-infection modifications (10).

Little is currently understood about the mechanisms how *T. annulata* manipulates host signaling pathways post-invasion, contributing to uncontrolled proliferation. In *Plasmodium* and *Toxoplasma*, the release of effector proteins before, during, and after invasion is critical to host cell entry and manipulation (11, 12). In contrast, the release of microsphere and rhoptry proteins by *Theileria* appears to occur only after complete internalization of infectious sporozoites (13). Interestingly, and unlike other apicomplexan zoites, the *Theileria* sporozoite does not need to reorient itself to bring its apical pole into close contact with the host cell membrane but enters the host cell in any direction through a progressive so called “circumferential zippering mechanism” (14). After the invasion process, *Theileria* parasites only briefly remain inside a parasitophorous vacuole membrane (PVM), from which they escape within minutes of infection to lie free in the host cell cytoplasm (15). Immediately after exiting the PVM, the parasite associates with host cell microtubules (13, 16). The unique positioning of *Theileria* parasites inside the host cell cytoplasm, not enclosed by a PVM, facilitates the recruitment, manipulation, and hijacking of host cell proteins such as EB1 and IKK directly on their membrane surface (4, 17, 18), distinguishing them from the other apicomplexan parasites. Approximately three to four days after leukocyte invasion by tick-born sporozoites, the developed multinucleated schizont triggers uncontrolled clonal proliferation and host cell immortalization (19, 20), with the unique feature that each subsequent host cell division results in an equal distribution of the *Theileria* schizont between the two daughter cells (16, 21, 22). The transformed parasitized host leukocytes eventually start to spread throughout the lymphoid system and the rest of the body (23, 24). Within the parasite’s life cycle, the schizont stage plays a central role in the pathology associated with malignant *T. annulata*, setting it apart from other *Theileria* species such as non-malignant *T. orientalis* strains. In *T. orientalis* the intra-erythrocytic piroplasm stage is responsible for the pathology, and no host cell transformation occurs (25, 26).

Infected cells in malignant theileriosis exhibit significant changes in gene expression profiles, including the persistent activation of the phosphatidylinositol 3-kinase (PI3-K) pathway (27), upregulation of c-Jun NH2-terminal kinase (JNK) (28, 29), increased c-Myc expression (30), and suppression of p53 activity (31, 32). The schizont hijacks the I_K_B kinase (IKK) signalosome on the schizont membrane, activating the NF-_K_B pathway and influencing anti-apoptotic gene expression (17). Host manipulation induced by *T. annulata* is likely to involve the secretion of various effector proteins, which affect host signaling pathways and contribute to the uncontrolled proliferation, invasiveness, and resistance to apoptosis (33). Despite extensive studies on parasite-induced changes to the host cell, only a limited number of exported proteins, including TaPIN1 (34), TashAT2 (35), and Ta9 (36) have been identified. These effector proteins are suggested to be associated with specific host signaling changes that potentially contribute to the uncontrolled and invasive cancer-like host cell behavior (36–40). However, a comprehensive understanding of the essential exportome responsible for the malignant alteration of the host cell is still lacking.

We employed an unbiased approach, using TurboID-based proximity labelling (41) of *T. annulata*-infected TaC12 cells, aiming to identify secreted effector proteins in the host cell cytoplasm and nucleus. This strategy revealed a novel gene family named Nuclear Intrinsically Disordered Proteins (NIDP1 - 4), alongside members of the Tash and Ta9 protein families. Antibodies against NIDP1 - 4 confirmed their nuclear localization. Remarkably, this gene family was absent in the non-transforming parasite *T. orientalis*. Detailed analysis of NIDP2 showed a biphasic localization pattern of the protein, accumulating in the host cell nucleus during interphase and associating with a protein complex at the schizont surface during mitosis. Within the nucleus, NIDP2 localized to the host chromatin and interacted with the tumor suppressor STAG2, shedding light on potential mechanisms underlying host cell transformation of cancer-related pathways. Our study provides a novel perspective on the shared characteristics of *T. annulata* effector proteins, most strikingly their predicted intrinsic disorderedness, offering insights into their role in host cell manipulation.

## RESULTS

### TurboID-based proximity ligation in the host cell leads to the identification of secreted Theileria annulata proteins

We performed TurboID-based proximity labelling in the host cell cytoplasm and nucleus, with the aim of identifying new effector proteins secreted by the parasite, which might be involved in host cell transformation. For this, the promiscuous biotin ligase TurboID (41) was fused either to a nuclear localization signal (NLS) or a nuclear export signal (NES) sequence and expressed in the *T. annulata*-infected TaC12 macrophage cell line (**Fig. 1A**). By adding biotin to the cell culture media, proteins near the biotin ligase enzyme were biotinylated, affinity purified with streptavidin and subsequently identified by mass spectrometry (LCM-MS/MS). Immunofluorescence analysis (IFA) confirmed the correct localization of both fusion proteins in the continuously parasitized cell line TaC12 (**Fig. 1A**, **Fig. S1A**). Non-biotin controls were used for comparison and analyzed in parallel with three biological replicates for each construct. As an initial step in identifying new effector candidates, the detected peptide count in LC-MS/MS of the control was compared with the biotinylated samples. Based on these results, candidate proteins were prioritized according to the following criteria: The protein must occur in at least two of three replicates and contain a predicted signal peptide (SP) according to predictions of the SignalP 4.1 algorithm used with SignalP 3.0 sensitivity (42). These stringent criteria lead to nine candidate proteins predicted to be exported to the host nucleus, and four candidate proteins in the host cytosol (**Fig. 1B**). The full set of 179 *T. annulata* proteins identified with at least one peptide in at least one replicate is provided in **Supplemental Table S1**. We then searched for orthologues in the non-transformative species *T. orientalis*) (43), hypothesizing the absence of orthologous proteins in case of importance for transformation. A BLAST analysis of the nine proteins identified in the nuclear fraction found that two proteins TA09456 and TA17425, to share sequence identities with *T. orientalis* proteins TOT_0300000391 and TOT_030000030, respectively. In the cytosolic fraction, TA09615 and TA03615 were found to share identities with TOT_010001127 and TOT_030000583, respectively. TA03615 has been previously described as associated with the parasite membrane (16). TA16090 was identified in both nuclear and cytosolic fractions, and while no clear orthologue in *T. orientalis* is detected, some similarity (29% identity) is found in the N-terminal part of the protein with the *T. orientalis* protein TOT_010000916.

**Figure 1.**
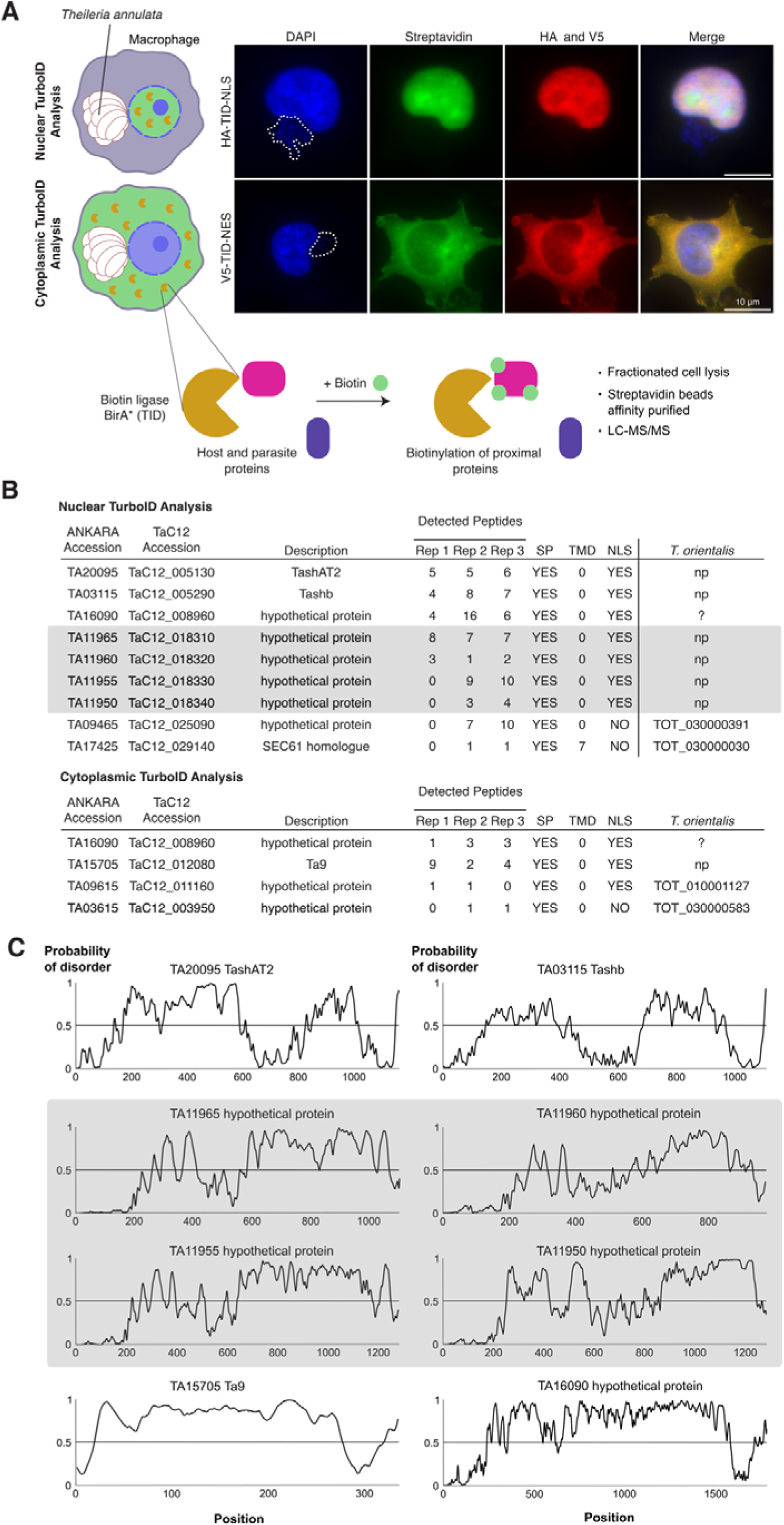
TurboID identifies proteins secreted by *T. annulata* into the host cell. **(A)** Schematic representation of TurboID (TID) approach in *Theileria annulata* TaC12 cells (infected macrophages). The biotin ligase TurboID was targeted to the host cell nucleus and cytoplasm. Upon addition of biotin for 3 h, biotinylated host and parasite proteins were fractionated and analyzed by mass spectrometry. To verify the correct localization and activity, HA-TID-NLS-construct transduced (host nuclear TID) and V5-TID-NES construct transduced (host cytoplasmic TID) TaC12 cells were grown in presence of biotin, fixed and analyzed by immunofluorescence assay using anti-HA or anti-V5 antibodies with additional staining of biotinylated proteins by FITC-conjugated streptavidin. The host cell nuclei and parasite schizont nuclei (indicated by dotted line) are labelled with DAPI. See also **Figure S1**. **(B)** Mass spectrometry results of three biological replicates from nuclear and cytoplasmic TID experiments with peptide counts of identified *T. annulata* proteins. Shown are proteins identified at least two times with a predicted signal peptide (SP) or transmembrane domain (TMD). See also **Table S1** for entire list. Highlighted in grey is a newly identified protein family with absent orthologues in non-transformative *T. orientialis*. **(C)** Protein disorder score (IUPred3) of all identified proteins with no orthologues in *T. orientalis*. NLS, nuclear localization sequence; np, not present.

Among the proteins in the host cell nuclear fraction with no orthologues in *T. orientalis*, we identified TashAT2 (TA20095) and Tashb (TA03115) (Fig. 1B). TashAT2 and Tashb are both members of the large Tash gene family clustered in tandem repeats on chromosome 1 (44) (**Fig. S1D**). We raised an antibody against the so far uncharacterized Tashb protein and confirmed that Tashb is targeted to the host cell nucleus of schizont-infected cells (**Fig. S1B**). TashAT2 has been previously identified as a secreted effector protein located inside the host cell nucleus of *T. annulata* D7 and TBL20 cell lines (35). We confirmed the nuclear localization of TashAT2 in TaC12 cells by IFA (**Fig. S1C**). Ta9 (TA15705) is the protein detected in the host cytoplasm with no orthologues in *T. orientalis* (Fig. 1B). Ta9 is a member of the Ta9 gene family (45, 46). We confirmed the anticipated localization in the host cytoplasm in TaC12 cells by IFA (**Fig. S1E**). Ta9 has previously been shown to be secreted into the host cell in the *T. annulata*-infected cell line Thei and is suggested to be involved in AP-1 transcription factor (TF) activation when overexpressed in embryonic kidney cells (36).

In the nucleus we identified four additional, so far uncharacterized proteins (Fig. 1B). Subsequently, we conducted a bioinformatic search for commonalities among the identified exported proteins and discovered that all those lacking an orthologue in *T. orientalis* exhibit a high intrinsically disordered protein structure, as predicted by IUPred3 (Fig. 1C) (47). Predictions by flDPnn (48) provided similar results (data not shown).

### Identification of a novel protein family of secreted proteins

The four so far undescribed *T. annulata* proteins in the nuclear fraction, TA11950, TA11955, TA11960 and TA11965, cluster in tandem repeats on chromosome 2 (Fig. 2A). Within this locus, the four proteins are flanked by an array of six considerably shorter genes (TA11945 to TA11900). Notably, TA11945 bears the closest homology to TOT_020000195 in *T. orientalis* as suggested by a phylogenetic analysis of the 10 genes of *T. annulata* and the four genes of *T. orientalis* present in the same locus on chromosome 2. The resulting tree showed a clustering of TA11950, TA11955, TA11960 and TA11965 on a separate branch distinct to the six smaller proteins in *T. annulata* and the four proteins present in *T. orientalis* (Fig. 2A; **Fig. S2A**).

**Figure 2.**
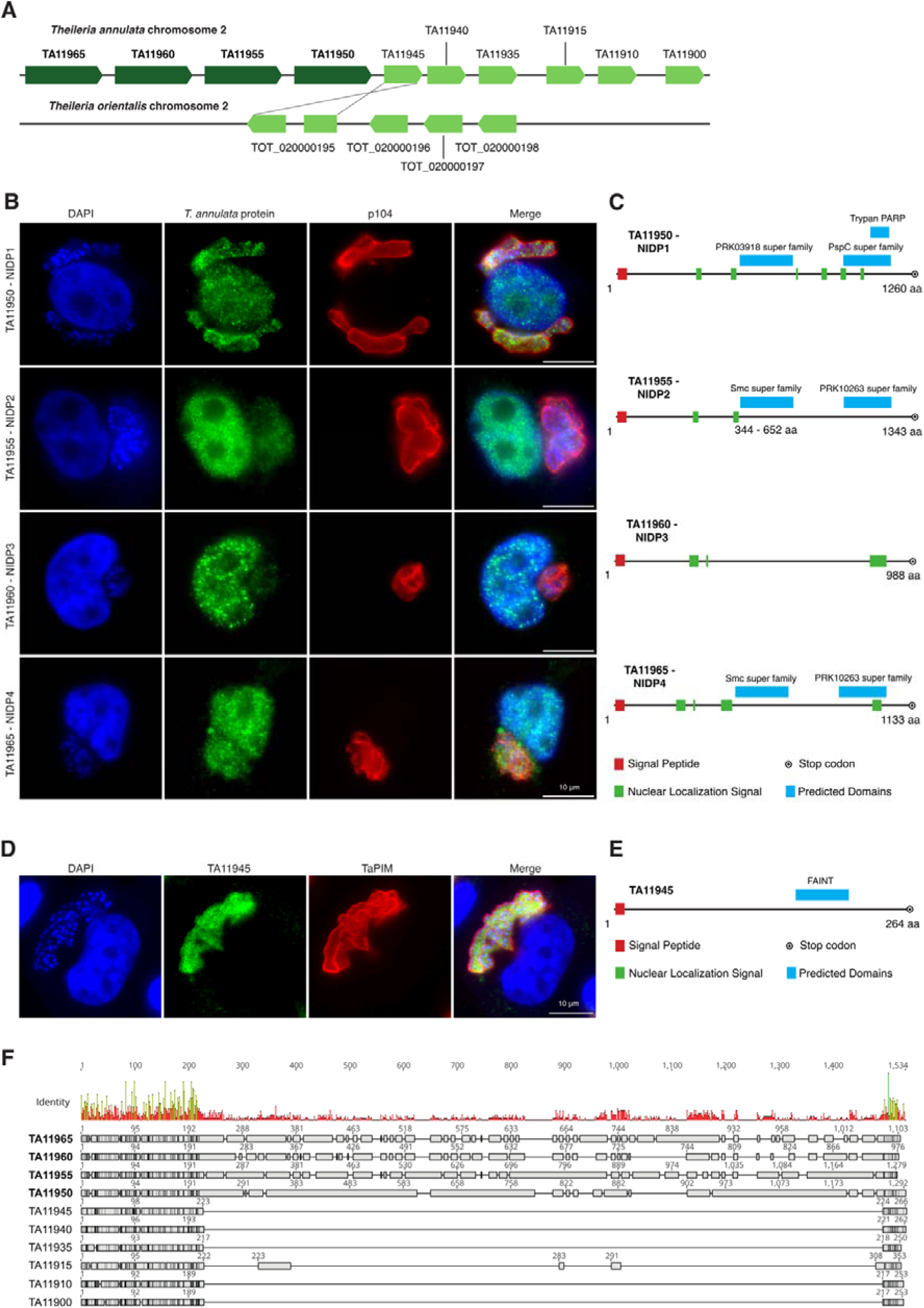
Characterization of a novel secreted Nuclear Intrinsically Disordered Protein (NIDP) family of *T. annulata*. **(A)** Schematic of protein locus on chromosome 2 of *T. annulata* and *T. orientalis*, respectively. A phylogenetic analysis suggests that TA11945 is orthologues to TOT020000195. See also **Figure S2A. (B)** TaC12 cells stained with α-TA11950 (NIDP1), TA11955 (NIDP2), TA11960 (NIDP3) and TA11965 (NIDP4), and α-p104 (schizont membrane) confirm the translocation of all four proteins into the host cell nucleus. Host cell nuclei and parasite nuclei are labelled with DAPI. **(C)** Schematic representation of proteins NIDP1 - 4 with predicted domains highlighted. **(D)** TaC12 cells stained with α-TA11945 and α-TaPIM (schizont membrane). Host cell nucleus and parasite nuclei are labelled with DAPI. **(E)** Schematic representation of protein TA11945 with predicted FAINT domain highlighted. **(F)** Alignment of *T. annulata* proteins NIDP1 - 4 with the other six members of the repetitive protein locus on chromosome 2.

Next, we raised antibodies against TA11950, TA11955, TA11960 and TA11965, and confirmed their localization inside the host nucleus of the TaC12 cell line as well as inside the parasite schizont syncytium (Fig. 2B). Notably, no staining was observed in non-infected control cells or with pre-immune serum (**Fig. S2B**). Subsequently we named the newly identified proteins *Theileria annulata* <u>N</u>uclear Intrinsically <u>D</u>isordered <u>P</u>rotein 1 (NIDP1; TA11950), NIDP2 (TA11955), NIDP3 (TA11960) and NIDP4 (TA11965). All four proteins contain a predicted signal peptide (SP) and predicted nuclear localization signals (NLS) (Fig. 2C). A domain search using the CDD/SPARCLE software (49) predicted the following conserved domains: For NIDP1, a homology to the PRK03918 super family (E-value: 9.77e-07) and a similarity to the trypan PARP region (E-value: 2.63e-10) and PspC superfamily (E-value: 5.00e-04) was found. The PRK03918 super family, or more precisely DNA double strand break repair ATPase Rad50, is a protein which is part of the structural maintenance of chromosome (SMC) protein family. Rad50 is also part of the MRN complex (MRN: complex consisting of MRE11, Rad50 and NBS1), which is implicated in DNA double strand break repair (DBS), break recognition, DNA end processing and functions as a signal for cell cycle arrest (50). NIDP2 and NIDP4, both contain a predicted domain for the conserved SMC super family (E-value NIDP2: 2.68e-09; NIDP4: 2.52e-04) as well as the PRK10263 super family (E-value NIDP2: 1.42e-03; NIDP4: 1.40e-05) (Fig. 2C). SMC proteins are necessary for chromosome condensation before mitosis and during mitosis in sister chromosome resolution as well as siter chromatid cohesion. SMCs are also involved in DNA repair after mitosis and in the regulation of gene expression (51, 52). For NIDP3, no conserved domains were predicted. Taken together, the domain predictions suggest a potential involvement of three of the proteins in this family in regulation of gene expression and chromosome maintenance.

We also raised an antibody against the adjacent protein TA11945 and detected this protein only in the parasite schizont and not within the host cell (Fig. 2D). No staining was observed in non-infected control cells or with pre-immune serum on TaC12 cells (**Fig. S2C**). Unlike TaNIDP1 - 4, TA11945 harbors a FAINT domain (frequently associated in *Theileria;* <u>(Pain,</u> <u>2005)</u> of unknown function and no NLS (Fig. 2E).

To gain further insights into the overall protein structure of the arrayed protein family, we aligned the ten members of the protein family and analyzed their amino acid similarity. Surprisingly, all members showed high sequence identities at the N-and C-terminal end (**Fig. S2E**). We used alphaFold2 predictions (53) of NIDP2 and TA11945 to further highlight the structural similarities between both proteins. Interestingly, whereas the N- and C-termini of NIDP2 overlap with TA11945, the center of NIDP2 – (TA11955_252-1250_) – has largely expanded (**Fig. S2D, E**), as has NIDP1, NIDP3 and NIDP4 (Fig. 2F). Large parts of these protein expansions appear highly disordered (Fig. 1C).

### NIDP2 associates with host cell chromatin and is also found in the cytoplasm

NIDP2 is a highly disordered protein except for the N- and C-termini and few alpha helices which comprise the predicted SMC domain (Fig. 3A). Western blotting confirms that the protein is mainly localized in the host nucleus, with smaller amounts also detected in the cytoplasmic fraction (Fig. 3B). Although the predicted size of NIDP2 is 150.8 kDa, it is resolved with a higher apparent molecular weight of approximately 180 kDa by SDS-PAGE and Western blotting. Our IFA analyses indicate that proteins of the NIDP family might be expressed to some extent at the schizont surface (Fig. 2B). To investigate how NIDP2 interacts with the schizont membrane, we performed a Triton X-114 extraction and phase separation of TaC12 whole cell lysates to separate hydrophilic from amphiphilic membrane proteins that become enriched in the detergent phase (54). We detected NIDP2 in both the aqueous phase and pellet fraction, in contrast to TaSP (TA17315) which, as a transmembrane protein, is enriched in the detergent fraction (55, 56) (Fig. 3C). This confirmed the prediction that NIDP2 contains no transmembrane domain, nor any GPI anchor. To determine whether NIDP2 in the pellet fraction is insoluble or chromatin-associated, we fractionated the cells into cytoplasmic, nuclear, chromatin-bound and pellet fractions (57). Immunoblotting revealed NIDP2 in the cytosolic, nuclear and chromatin bound fraction, while the positive control Lamin B1 was almost exclusively found in the chromatin bound fraction. This may indicate that NIDP2 can associate with the host chromatin (Fig. 3D). Of note, a slight double band is observable in the nuclear fraction and in the chromatin bound fraction (Fig. 3B, D) possibly indicating post-translational modification (PTM) of NIDP2 in the host nucleus, predominantly when associated with the chromatin.

**Figure 3.**
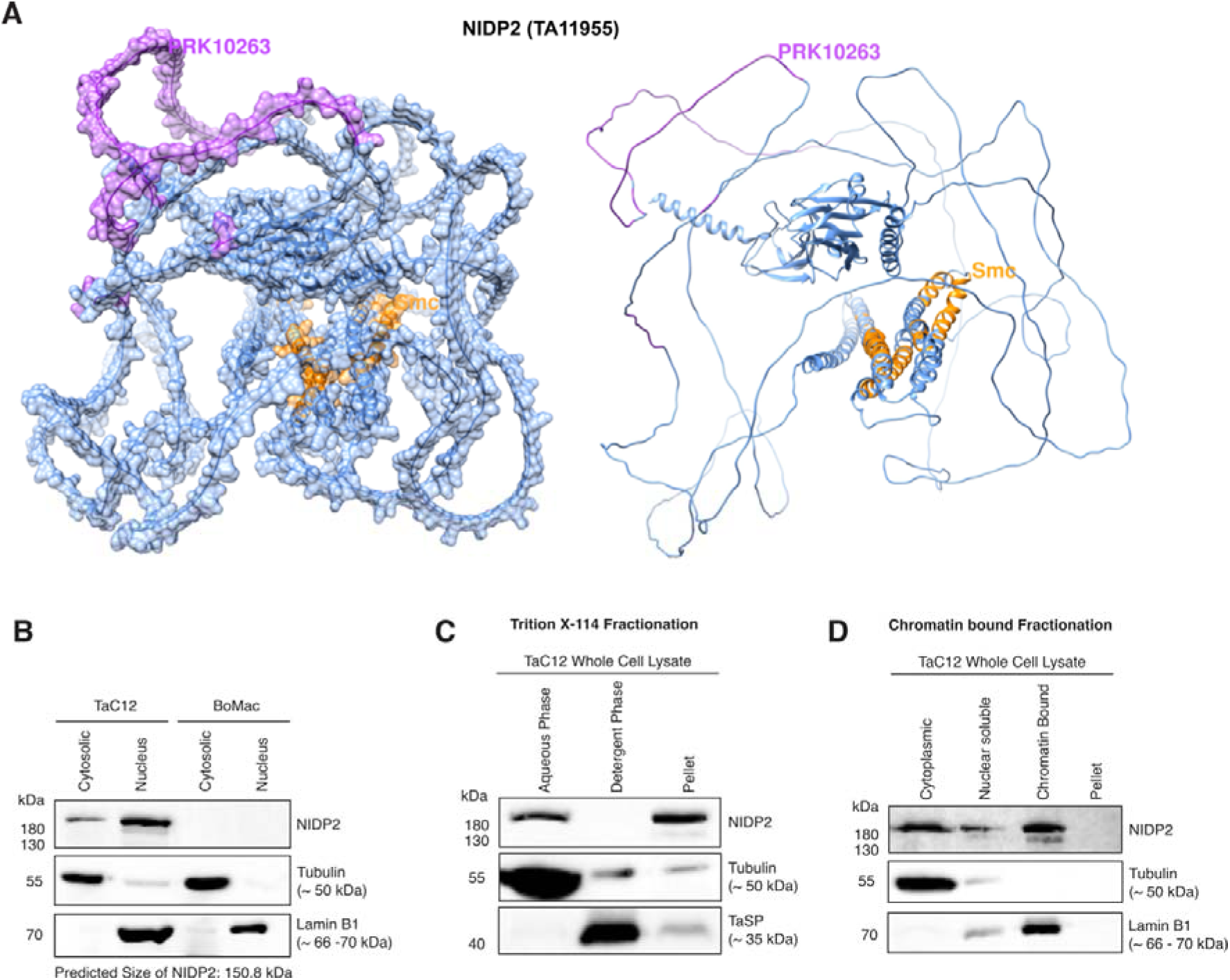
Characterization of Nuclear Intrinsically Disordered Protein 2 (NIDP2) **(A)** Predicted structure and domains of NIDP2 using alphaFold2. **(B)** Cytoplasmic and nuclear protein fractions of TaC12 and BoMac cells showed a stronger signal in the nuclear compared to the cytosolic fraction for NIDP2 >180 kDa. **(C)** Tac12 cells were fractioned with Triton X-114. NIDP2 could be detected in the aqueous phase and the pellet phase. Tubulin was used as control for the aqueous phase while TaSP (TA17315), a *Theileria* surface parasite protein containing a transmembrane domain, functioned as control for the detergent phase. **(D)** Cytoplasmic, nuclear soluble, chromatin bound and pellet fraction of TaC12 cells were analyzed by SDS-PAGE. NIDP2 was detected in the cytoplasmic, nuclear soluble and chromatin bound fraction. Tubulin was used as control for the cytoplasmic fraction, and laminB1 for the chromatin bound fraction.

### NIDP2 localizes to the schizont membrane via the CLASP1/CD2AP/EB1-complex in a cell-cycle dependent manner

To further explore the potential function of the NIDP protein family, we decided to investigate their localization throughout the cell cycle of the host cell. Strikingly, NIDP2, but not NIDP1, NIDP3 or NIDP4, localized exclusively to the parasite membrane during host cell mitosis, while all four family members are detected in the host nucleus during interphase (Fig. 4A, **Fig. S3A**). The biphasic localization of NIDP2 suggests a tightly regulated interaction with specific host and potentially other parasite proteins in two distinct compartments in a spatial and temporal manner. As the host cell enters mitosis and the host nuclear membrane breaks apart, NIDP2 colocalizes with the *T. annulata* protein p104 (18, 58) on the schizont surface (Fig. 4A, **Fig. S3A**). No residual staining of NIDP2 was observed close to or around condensed chromosomes until the host cell enters telophase/G1. The parasite membrane protein p104 has been shown to interact with host end binding protein 1 (EB1) and CLIP-170-associating protein 1 (CLASP1) on the parasite surface. EB1 is an important regulator of MT dynamics and CLASP1 is a microtubule stabilizing protein (16, 18). In addition to EB1 and CLASP1, the CD2-associated protein (CD2AP) can be found as part of this larger protein complex on the schizont surface (59). Notably, CLASP1 and CD2AP are present on the parasite surface during the whole cell cycle of the host cell (16, 59). To further investigate the potential interaction of NIDP2 with the CLASP1/CD2AP/EB1-complex, we successfully coimmunoprecipitated p104 and CLASP1 together with NIDP2 in TaC12 cells (Fig. 4B). In addition, we engineered CD2AP-TurboID and CLASP1-TurboID constructs that target the schizont membrane throughout the cell cycle, and stably expressed the fusion proteins in TaC12 cells (Fig. 4C, **Fig. S3B**). After subcellular protein fractionation, affinity purified biotinylated proteins were analyzed by mass spectrometry in triplicates and the results were categorized as described before. NIDP2 was detected in both schizont-surface TurboID analyses (Fig. 4D, E). Our data therefore suggest that NIDP2 is a member of the EB1/CD2AP/CLASP1-complex. Unlike other parasite protein members of this complex such as p104 and MISHIP (59), NIDP2 translocates to the host nucleus during interphase. The interaction with the EB1/CD2AP/CLASP1 complex on the schizont surface appears to be transient and is not a result of NIDP2 integration into the parasite membrane (Fig. 3C).

**Figure 4.**
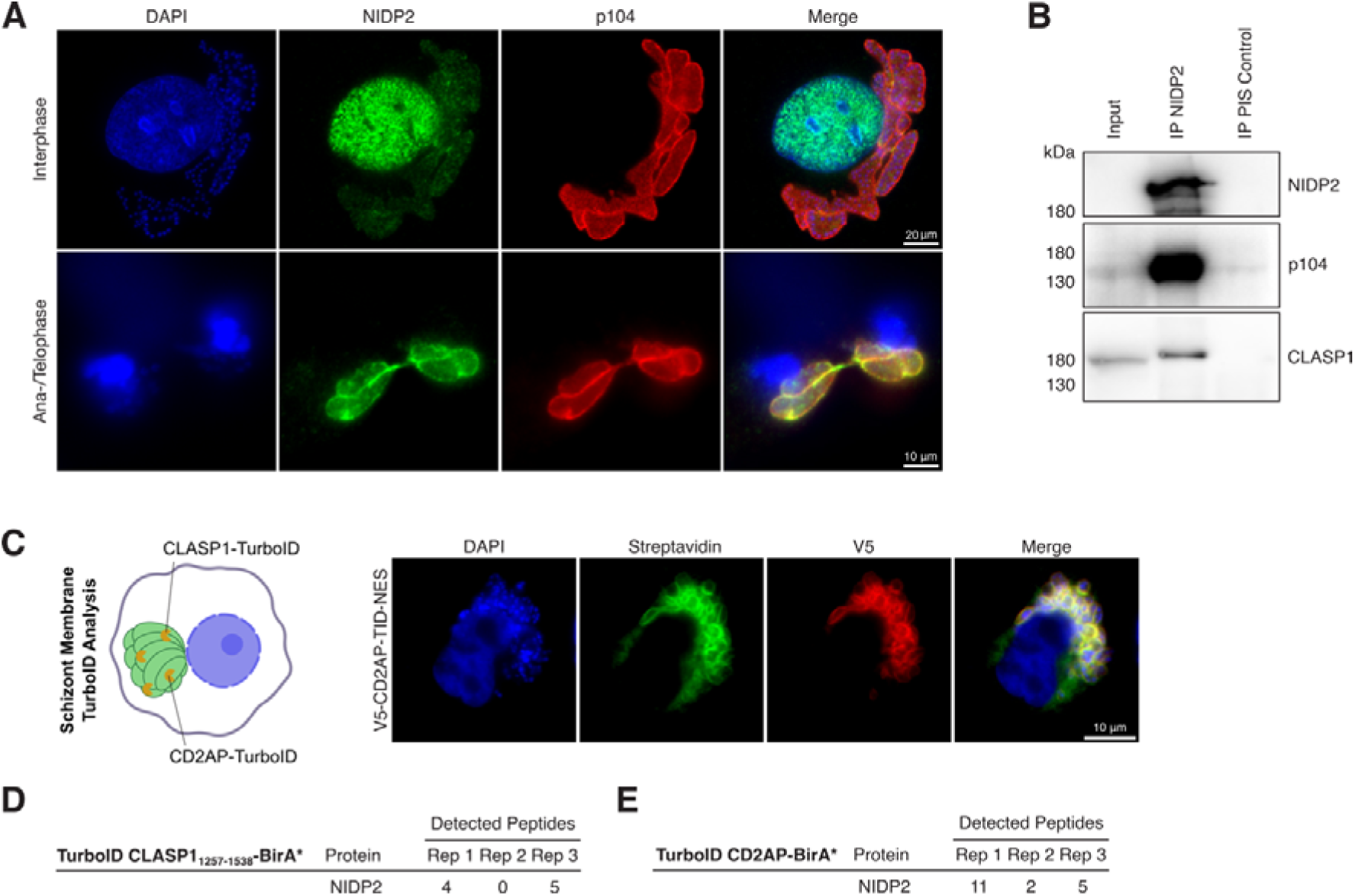
Cell cycle dependent localization of NIDP2 in host nucleus and CLASP1-CD2AP-EB1-p104-TaMISHIP parasite membrane complex. **(A)** TaC12 cells stained with α-NIDP2 in interphase (upper panel) and ana-/ telophase (lower panel). The schizont surface is stained with α-p104, and host and parasite nuclei with DAPI. Note the colocalization of p104 and NIDP2 during mitosis. See also **Figure S4**. **(B)** Western blot analysis of NIDP2 immunoprecipitation (IP) from TaC12 cells (15% of total) blotted with primary antibodies as indicated. Preimmune serum (PIS) from the same rabbit served as control. Representative of three experiments with similar outcomes. **(C)** Schematic representation of TurboID analysis with TurboID-CD2AP and TurboID-CLASP1_1256−1538_ fusion proteins targeted to the parasite surface in TaC12 cells. Biotin-treated, V5-TID-NES-CD2AP-construct-transduced TaC12 cells show specific biotinylation of the parasite membrane (FITC-conjugated streptavidin; green). Host cell nucleus and parasite nuclei are labelled with DAPI. **(D)** CLASP1-associated NIDP2 peptides identified by LC-MS/MS analysis in three biological replicates. **(E)** CD2AP-associated NIDP2 peptides identified by LC-MS/MS analysis in three biological replicates. For clarity, only NIDP2 peptides are shown.

### *In vivo* cross-linking of NIDP2 identifies proteins involved in cancer as potential host nuclear binding partners

To gain insights into the role of NIDP2 in the host nucleus of *T. annulata* infected macrophages, we performed *in vivo* cross-linking and immunoprecipitated NIDP2 protein complexes in three biological replicates from nuclear extracts of TaC12 cells. As controls, we immunoprecipitated with rabbit preimmune serum from nuclear extracts of TaC12 cells and with α-NIDP2 from nuclear extracts of non-infected BoMac cell lysates (Fig. 5A). Protein complexes of all replicates and controls were analyzed by mass spectrometry (LC-MS/MS). Only proteins, which were identified in α-NIDP2 pulldown assays from TaC12 cells and not in the controls, were considered potential interactors of NIDP2 within the host nucleus. Aside NIDP2, we did not pulldown any other members of the NIDP family.

**Figure 5.**
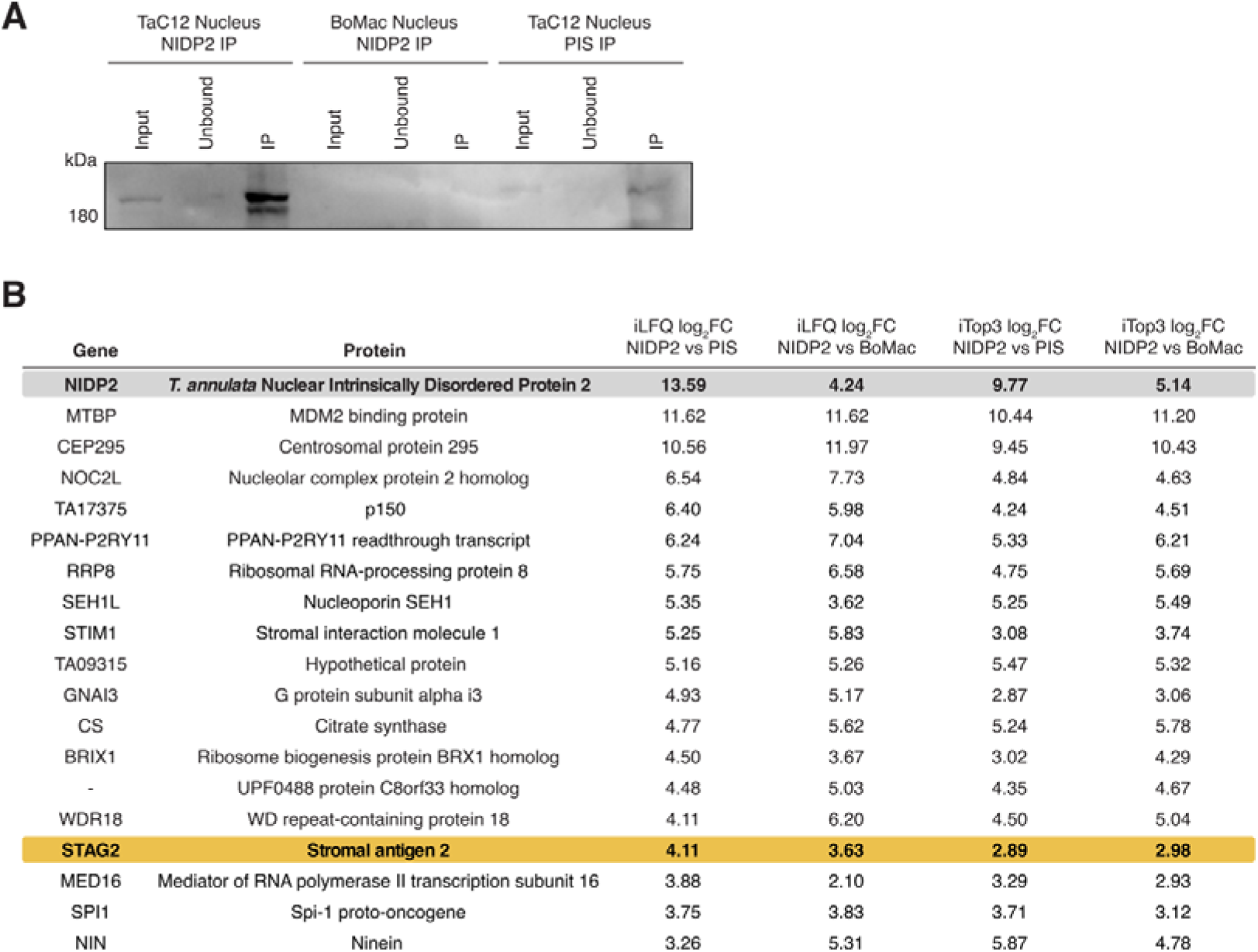
Identification of interaction partners of NIDP2 in the host nucleus. **(A)** Western blot analysis of nuclear fractions of TaC12 (infected) and BoMac (uninfected) cells immunoprecipitated (IP) with α-NIDP2; TaC12 cells were additionally probed with pre-immune serum (PIS) from the same rabbit. The Western blot was probed with α-NIDP2 as primary antibody. **(B)** NIDP2 associated parasite and host proteins identified by LCM-MS/MS analyses are shown. The iLFQ log_2_ fold change (FC) was calculated between three biological replicates of the NIDP2-IP of TaC12 cells and two biological replicates of the PIS-IP. The iLFQ log_2_ FC NIDP2 vs BoMac was calculated between three biological replicates of NIDP2-IP of TaC12 cells and two biological replicates of NIDP2-IP of BoMac cells. The iTOP3 log_2_ FC values were calculated in the same way.

As potential host binding proteins, we identified multiple proteins implicated in cancers. Two proteins are implicated in the regulation of p53: Mouse double minute 2 (MDM2) binding protein (MTBP) (60) and Nucleolar complex protein 2 homolog (NOC2L) (61) (Fig. 5B). Importantly, we also identified Stromal Antigen 2 (STAG2), which serves as tumor suppressor and an accessory protein of cohesin complexes (62). Cohesin, a protein complex associated with Structural Maintenance of Chromosomes (SMCs), plays a critical role in sister chromatid cohesion, chromosome condensation, DNA repair, 3D genome organization and gene expression, and is among the most commonly mutated protein complexes in cancer (63, 64).

### NIDP2 interacts with STAG2 in the nucleus of the host cell

Given the predicted SMC domain for NIDP2 (Fig. 2C, **3A**) and considering that STAG2 is frequently mutated in various cancers (65), we decided to investigate the NIDP2-STAG2 interaction further by IFA of both proteins in TaC12 cells. This revealed a high level of co-localization of both proteins within the host cell nucleus, while no such co-localization was found when NIDP2 was localized on the parasite surface during host cell division (Fig. 6A, B).

**Figure 6.**
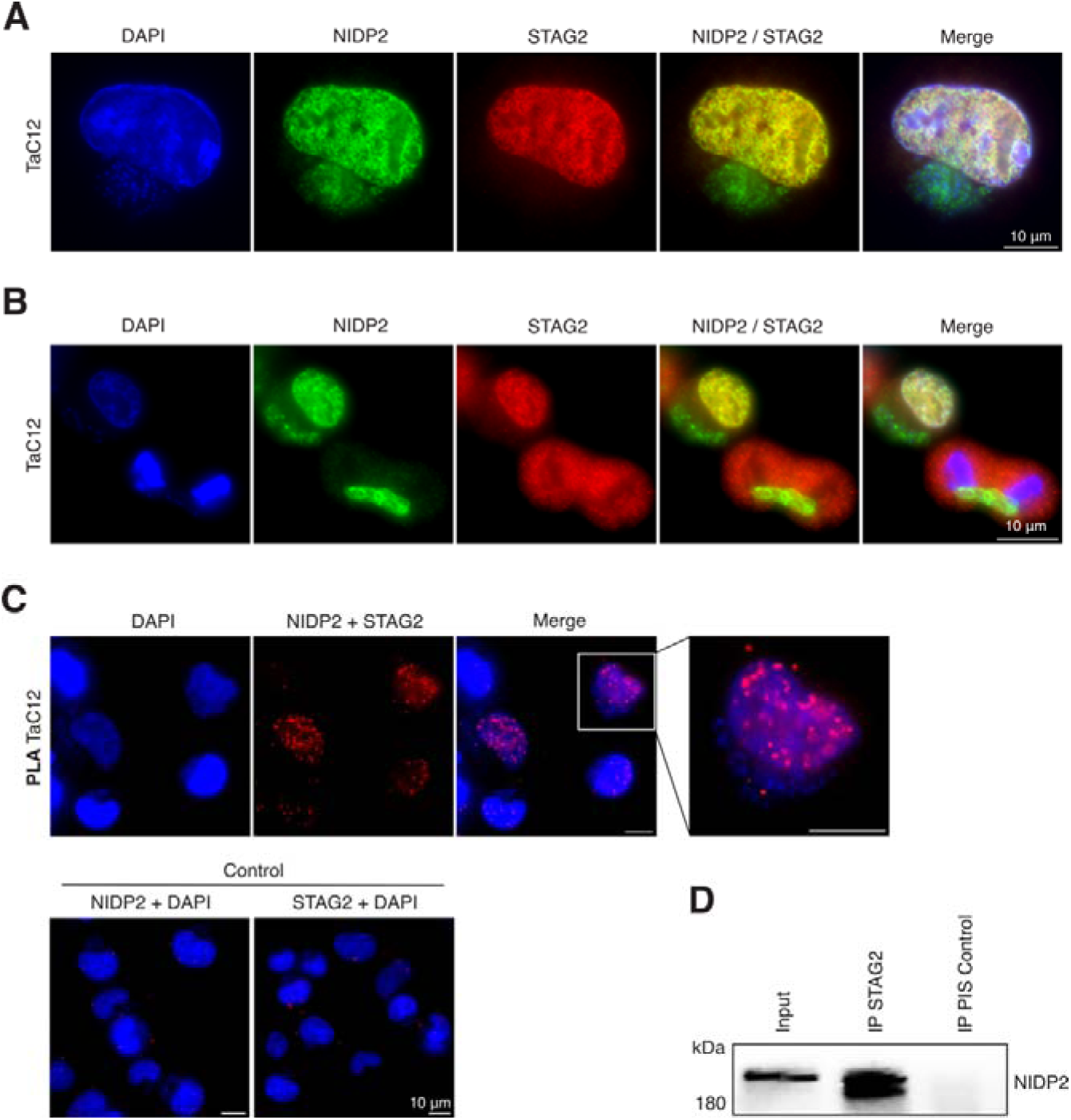
NIDP2 interacts with STAG2 in the host cell nucleus. **(A)** TAC12 cells stained with α-NIDP2 and α-STAG2 show colocalization of both proteins inside of the host nucleus, but not on the parasite surface **(B)**. Host and parasite nuclei are stained with DAPI. **(C)** Proximity ligation assay (PLA) of TaC12 cells shows a signal in host cell nuclei only in the presence of both α-NIDP2 and α-STAG2. **(D)** Western blot analysis of STAG2 immunoprecipitation (IP) from TaC12 cells (20% of total) blotted with α-NIDP2. Preimmune serum (PIS) from the same rabbit served as control. Representative of three independent experiments with similar outcomes.

To further corroborate this finding, we utilized a proximity ligation assay (PLA) that produces a fluorescent signal when two proteins are within 40 nm of each other. This assay revealed a signal for the NIDP2-STAG2 antibody-combination but no signal when both antibodies were applied alone (Fig. 6C), providing further evidence of the interaction of both proteins within the host cell nucleus. In line with this, we were also able to further confirm the interaction of STAG2 with NIDP2 by Western Blot analysis after STAG2 immunoprecipitation from TaC12 cells (Fig. 6D).

## DISCUSSION

Transforming *Theileria* species uniquely induce a cancer-like state in infected host cells, marked by a heightened proliferation, immortality, invasion, and metastasis. The specific parasite proteins and mechanisms driving these profound host cellular changes remain poorly understood, and, apart from bioinformatic predictions (34, 43, 46), an unbiased experiment to identify exported effector proteins has not been conducted. By employing a TurboID-based proximity labelling approach targeting different host compartments in *T. annulata-*infected macrophages, we identified several exported proteins and revealed a common feature among *Theileria* effector proteins, namely a high protein disorder score, indicative of intrinsically disordered proteins (IDPs). In addition to known exported proteins, a novel protein family, <u>N</u>uclear Intrinsically <u>D</u>isordered <u>P</u>roteins (NIDP) 1 – 4, clustered in tandem repeats on chromosome 2, was discovered. The identification of this new protein family, alongside members of the Ta9 and Tash families, underscores the significance of expanded protein families for transformative *T. annulata*. Our investigation into NIDP2 revealed an intriguing biphasic cell cycle-dependent localization, demonstrating interactions with the EB1/CD2AP/CLASP1-complex on the parasite membrane during mitosis and with the tumor suppressor STAG2 within the host cell nucleus during interphase.

In our approach, we focused on identifying parasite proteins exported to the host cell nucleus and cytoplasm. Alongside the newly identified NIDP1 - 4 protein family, we verified the presence of known exported proteins: Ta9 (TA15705) in the host cytoplasm and TashAT2 (TA20095) in the host nucleus of TaC12 cells (35, 36). Additionally, the previously uncharacterized protein Tashb (TA03115) was found in the host nucleus. These identifications highlight the suitability of our TurboID-based approach for the discovery of exported *Theileria* proteins. Notably, Ta9, TashAT2, Tashb, and NIDP1 – 4, all of which are absent from the genome of the non-transforming *T. orientalis* (43), belong to larger gene families characterized by tandem repeats and variable copy numbers. Genes critical for survival duplicate under selective pressure indicating adaptive evolution, resulting in copy number variation (66), possibly driven by their role in pathogenesis and invasiveness (67, 68). Examples from other protozoans, such as the *vsg* and *var* and MEDLE gene families in trypanosomes*, Plasmodium* and *Cryptosporidium*, respectively, emphasize this evolutionary mechanism (69–71). Expanded gene families such as the *T. annulata* NIDP family are strikingly absent from the non-transforming *T. orientalis.* Notably, NIDP1 - 4 are tandemly arranged in a family of 10 proteins, with only one non-exported protein identifiable as an orthologue to a *T. orientalis* protein. The *T. orientalis* genome also harbors only a single-copy Tash gene. It is noteworthy that the orthologue of this gene in *T. annulata*, known as Tasha, is not expressed during the transformative schizont stage of the parasite (43). Furthermore, the *T. annulata* Ta9 gene shares only weak homology with the signal peptide region and C-terminal region of a *T. orientalis* gene (43). While both species infect leukocytes and develop into multinucleated schizonts, *T. annulata* induces uncontrolled lymphoproliferation prior to merogony, a feature absent in *T. orientalis*. Ultimately, our findings support the concept that extended gene families play a crucial role in the transformative capacity of *T. annulata* and may reflect the intimate host-pathogen co-evolution driven by an arms race between the parasite and its host (72).

In addition to the role of tandem arrayed proteins in malignant *Theileria*, the significance of intrinsically disordered proteins (IDPs) or intrinsically disordered regions (IDRs) in exported proteins among apicomplexans remains largely unexplored. For instance, *Toxoplasma* has numerous exported dense granule proteins such as GRA24, GRA16 and TgIST, all of which are characterized by distinct disordered protein structures (73–76). The structural flexibility and lack of a well-defined three-dimensional structure may allow for the interaction with multiple host protein partners, potentially increasing functional complexity (12). The dynamic nature and rapid evolution of unstructured regions may also optimize the efficacy of the effectors in the parasite’s arsenal and potentially also influence trafficking across the parasitophorous membrane. Because IDRs lack a defined protein structure, they may facilitate a less energy-costly export of effector proteins across the parasitophorous vacuole membrane (PVM), as they do not require unfolding to pass through a membrane channel such as the PTEX complex in *Plasmodium* (77). Specifically, these *Toxoplasma* effector proteins traverse the PVM via interaction with the putative MYR1 translocon (78), and the presence of structured tags impedes translocation, leading to protein entrapment within the parasitophorous vacuole (79). However, unlike *Toxoplasma* and closely related *Plasmodium*, *Theileria* (like *Babesia*) lacks a PVM, residing freely in a single membrane syncytium within the host cell’s cytosol, and no orthologues of the MYR1 or PTEX translocon protein members have been identified for *Theileria* (4). This suggests that protein export in *Theileria* occurs through an unrelated mechanism. Protein disorder may not be a prerequisite for membrane translocation into the host cell in *Theileria*. This notion is supported by the protein TaPIN1, the only structured *T. annulata* protein identified so far exported into the host cell (34). Hence, the IDP signature may not be essential for export but might reflect a rapid and evolutionarily cost-effective process for expressing novel interactors with versatile functions. This phenomenon is exemplified in the NIDP protein family. These differences in protein export mechanisms highlight the diversity in strategies employed by apicomplexan parasites for interacting with and manipulating their host cells. A comparative analysis of exported effector proteins in apicomplexans may provide valuable insights into the evolutionary advantage behind the IDP/IDR signature (80, 81).

Our structural analysis of NIDP proteins, especially NIDP1 - 4, indicates extensive disordered expansions between the N- and C-terminal conserved regions of the arrayed protein family. As less defined protein structures are critical for the diverse functions of IDPs in key biological processes, such as signal transduction, transcriptional regulation, and cell cycle control (82), this feature may allow them to engage in promiscuous interactions with multiple protein binding partners. During interphase, NIDP2 localizes inside the host cell nucleus where stringent mass spectrometry analyses suggest multiple protein interaction partners including STAG2, mouse double minute 2 (MDM2) binding protein (MTBP), and nucleolar complex protein 2 homolog (NOC2L), the latter two both involved in p53 regulation. We successfully validated the interaction of NIDP2 and STAG2, a cohesion complex member and well-stablished cancer gene associated with various malignancies, including acute myeloid leukemia and bladder cancer (65). Notably, both p53 and MDM2 regulation has been previously shown to be altered in *Theileria* infected cells (31, 32). Unfortunately, attempts to confirm the interaction of NIDP2 with MTBP and NOC2L were inconclusive due to the unreliable performance of commercially available antibodies in the bovine background. There is evidence indicating that NIDP2 may undergo extensive additional posttranslational modifications when associated with chromatin, potentially contributing to its proper function in the host cell nucleus and hindering ectopic NIDP2 expression for functional interrogation.

Notably, during mitosis, NIDP2 relocates to the schizont membrane and interacts with the EB1/CD2AP/CLASP1 complex, which is involved in microtubule interaction and further interaction with parasite proteins p104 and TaMISHIP (4, 59). This raises intriguing questions about NIDP2’s dual function on the parasite surface and in the host cell nucleus, as well as its potential role in microtubule binding during mitosis. While attempts to ectopically express truncated forms of NIDP2 in bovine cells were unsuccessful, further work is needed to determine the function of NIDP2 in the nucleus, which regions of the protein interact with the EB1/CD2AP/CLASP1 complex, and whether NIDP2 interacts with additional proteins, as suggested by our mass spectrometry data set.

In conclusion, we identified a set of exported proteins characterized by a predicted high protein disorder score, notably the newly identified Nuclear Intrinsically Disordered Proteins (NIDP) family, alongside the established Ta9 and Tash protein families employing an unbiased TurboID-based proximity labelling approach. These findings challenge simplistic assumptions regarding the sole significance of protein disorder in facilitating protein export over the PVM in apicomplexan parasites. Instead, they study the existence of additional functional implications for the evolutionary development of protein disorder in apicomplexans. The detailed analysis of NIDP2’s biphasic cell cycle-dependent localization and interactions, including its association with the tumor suppressor and cohesion protein STAG2 and the EB1/CD2AP/CLASP1 membrane complex, sheds new light on previously unknown versatile dynamics of exported *Theileria* proteins. Collectively, these discoveries establish a foundation for further investigations into the molecular mechanisms governing *Theileria*-induced cancer-like host cell alterations.

## MATERIAL AND METHODS

### Maintenance of mammalian cell lines

TaC12 (*T. annulata* infected macrophage cell line), BoMacs (non-infected bovine macrophage cell line) (83) and HEK293T were cultured in flasks and well plates from TPP Techno Plastic Products AG, Switzerland. BoMacs and HEK293T were kept in DMEM at 37 °C and 5% CO_2_ atmosphere. TaC12 were cultured in L15 at 37 °C and 0% CO_2_. The culture media from Gibco were supplemented with 10% fetal calf serum (FCS; BioConcept, Allschwil, Switzerland; Cat. No. 2-01F10-I), 2 mM L-glutamine (BioConcept, Cat. No. 5-10K00-H), 100 IU/m penicillin and 100 µg/mL streptomycin (BioConcept, Cat. No. 4-01F00-H), 10 mM HEPES pH 7.2 (Merck, Darmstadt, Germany; CAS-No: 7365-45-9). TaC12 cells were washed with PBS and incubated with 1 mM PBS/EDTA (Merck) and the remaining cell lines were washed with PBS and incubated with 1x Trypsin-EDTA PBS (BioConcept, Cat. No. 5-51K00-H) until detachment. Cells were split with culture media as required.

### Expression constructs

All primers used in this study were ordered from Microsynth AG (Balgach, Switzerland), and constructs were verified prior to transfection by Sanger sequencing (Microsynth AG). Cloning was performed either with Gibson Assembly (New England Biolabs, NEB, Ipswich, MA) or In-Fusion HD cloning kit (Takara Bio, San Jose, CA) following the manufacturer’s instructions. Annealing temperatures were calculated using NEB Tm Calculator version 1.13.1 (https://tmcalculator.neb.com/#!/main). Plasmids were transformed into NEB 5-alpha Competent *E. coli*, and lentiviral constructs into Endura cells (BioCat, Heidelberg, Germany). The following plasmids were purchased from addgene (Watertown, MA, USA): pLenti CMV GFP Puro (658-5; 174488), second-generation packaging vector (psPAX2; 12260), VSV-G coat envelope vector (pMD2.G; 12259; Tronolab), V5-TurboID-NES_pCDNA3 (107169), 3xHA-TurboID-NLS_pCDNA3 (107171). For the lentiviral NLS-TurboID construct (lab number p1141) the BirA* region was amplified by PCR using primers F77 and F78 from plasmid 107171 and cloned into plasmid 174488, which was linearized beforehand with BamH1 and Sal1, to remove the EGFP region. For the lentiviral NES-TurboID construct (p1140), the BirA* region was amplified by PCR using primers F71 and F72 from plasmid 107169 and cloned into plasmid 174488, which was linearized as described above. For the lentiviral CD2AP-TurboID construct (p1138), the CD2AP gene was amplified from plasmid p869 as described earlier (59) with primers F11 and F12 and subsequently cloned into a PCR linearized plasmid 107169 with primers F13 and F14 containing overhangs for the CD2AP gene as well as a GSGS linker between the CD2AP and BirA*. In a second step the CD2AP-GSGS-TurboID fragment was amplified by PCR by primers F71 and F72 and cloned into plasmid 174488 which was linearized beforehand with BamH1 and Sal1. For the lentiviral CLASP1_1256−1538_-TurboID construct (p1139) the CLASP1_1256−1538_ region was amplified from plasmid p777 as described earlier (16) with primers F31 and F32 and subsequently cloned into a PCR linearized plasmid 107169 with primers F44 and F45 containing an overhang for the CLASP1_1256−1538_ gene as well as a GSGS linker in between the CLASP1_1256−1538_ and the BirA* construct. In a second step the CLASP1_1256−1538_-GSGS-TurboID fragment was amplified by primers F71 and F72 and cloned into plasmid 174488 which was linearized beforehand with BamH1 and Sal1. For the lentiviral mNeoGreen control plasmid (p1188), a TY1-mNeoGreen fragment was amplified by PCR using primers F79 and F80 from pcDNA4TetOn-TA06380-TY-mNeonGreen containing overlapping regions for the linearized pLenti CMV GFP Puro (658–5) (174488) plasmid which was linearized beforehand with BamH1 and Sal1 to replace the EGFP region. In a second step, a KOZAK sequence was added in front of the TY1 region with the Q5 Site-Directed Mutagenesis Kit (NEB) using primers F167 and F168. To generate GST fusion proteins, GST-TA11950_2203-2545_ was amplified from TaC12 gDNA with primers F220 and F221 and cloned into BamH1 restriction sites of the AF097411.1 expression vector pGST-parallel3. Accordingly, GST-TA11955_1642-1974_, GST-TA11960_1249-1617_ and GST-TA11965_1375-1698_ were amplified with primers F222 and F223, F224 and F225 and F226 and F227, respectively, and cloned into the pGST-parallel3 vector as described before.

### Lentiviral transduction and FACS sorting

HEK239T cells were used to produce lentiviruses which were transfected with FuGENE HD transfection reagent (Promega, Madison, WI; Cat. No. E2311) using a third-generation lentiviral transfer vector system. Briefly, pRRL-RSrII containing the gene of interest, packaging vector psPAX2 and envelope vector pMD2.G were transfected into HEK293T cells in a 5:3:2 ratio. Twenty-four hours post transfection media was replaced and lentiviral-particle-containing media was harvested 48 h and 72 h post-transfection. A 45 µm filter membrane was used prior to transduction of 2 x 10^5^ TaC12 wild type (WT) cells with 5 ml of the collected virus-containing media. TaC12 cells were transduced twice within 48 h with a recovery time of 24 h between both transduction steps. TaC12 cells expressing NES, NLS, CLASP1_1256−1538_- and CD2AP-TurboID constructs were sorted into a 96-well plate as single cells with the FACS sorter Aria III (BD Biosciences, San Jose, California, USA).

### Antibodies

The GST fusion proteins (GST-TA11950_2203-2545_, GST-TA11955_1642-1974_, GST-TA11960_1249-1617_ and GST-TA11965_1375-1698_) were overexpressed in BL21 Star *E. coli* (Invitrogen ThermoFisher Scientific, Waltham, MA). Proteins were purified by using glutathione Sepharose beads (GE Healthcare, Waukesha, WI), and subsequently sent to Eurogentec (Seraing, Belgium) for antibody production and purification in rabbits. Tashb (aa sequence: CQYVKSDSDNEENNND) and TA11945 (aa sequence: CEGVTESGELYSKSTY) were synthetized by Eurogentec and used for immunization in rats (Eurogentec, Seraing, Belgium).

### Immunofluorescence assays

Cells were seeded onto glass coverslips and incubated overnight and treated or remained non-treated prior to fixation with 4% PFA for 15 min at room temperature (RT) before washing with PBS and permeabilization in 0.2% Triton XC100 (diluted in PBS) for 10 min. Subsequentially cells were blocked in 10% FCS in PBS for 1 h at RT. Alternatively, cells were fixed with ice-cold methanol, washed twice with PBS and blocked in 10% FCS in PBS for 1 h at RT. Primary antibodies were diluted in 10% heatCinactivated FCS in PBS and put directly onto the cells for 1 h at RT. After primary antibody staining, cells were washed five times in PBS and secondary antibodies were diluted in 10% heatCinactivated FCS in PBS and incubated for 1 h at RT. DNA was stained using DAPI (Invitrogen), and samples were mounted onto slides by using mounting media (DAKO), if not mentioned otherwise. Freshly prepared samples were either analyzed on a DeltaVision Elite system (GE Healthcare) equipped with Olympus IXC70 inverted microscope and a CMOS camera, using a 100x Olympus Objective, and software from SoftWorx (Applied Precision) or an Eclipse 80i microscope (Nikon) equipped with a Hamamatsu Orca R2 camera using a 100x PlanApo objective (Nikon) and the OpenLab 5 software (Improvision). The PLA assay was performed according to the manufacturer’s instructions using the In Situ Detection Reagents Red (Catalogue Number: DUO92008, Merck, Darmstadt, Germany).

### TurboID to identify proteins interacting with CLASP1, CD2AP and located in the host cytoplasm and nucleus

TaC12 cells stably transduced with the described TurboID-fusion constructs NLS-TurboID, NES-TurboID, CD2AP-TurboID and CLASP1_1256−1538_-TurboID were incubated in media containing 500 µM Biotin (Serva) for 3 h (CLASP1_1256−1538_ is the minimal region for CLASP1 that still allows it to bind to the parasite membrane (16). As control the same cell lines were incubated for 3 h without Biotin. Cells were washed with PBS prior to detachment. Proteins were extracted using the NE-PER Kit (Thermo Scientific, Catalog Number 78833) for NLS-TurboID (nuclear fraction) and NES-TurboID (cytosolic fraction) and the Subcellular Protein Fractionation Kit (Thermo Scientific, Catalog Number 78840) for CD2AP-TurboID and CLASP1_1256−1538_-TurboID (membrane fractions). After protein extraction, lysates were incubated overnight at 4 °C with Pierce Streptavidin Magnetic Beads (Thermo Scientific, Catalog Number 88816) which were washed beforehand with RIPA Lysis Buffer (50 mM Tris pH 8, 150 mM NaCl, 0.1% SDS, 0.5% sodium deoxycholate, 1% Triton X-100, cComplete EDTA free protease inhibitor (Roche, Basel, Switzerland). Subsequently the magnetic beads were washed once with 1 x 1M KCL, 1 x 0.1 M Na_2_CO_3_, 1 x 2 M Urea in 10 mM Tris-HCl pH 8 and two times in RIPA Lysis Buffer. The supernatant was removed and beads were snap frozen prior to mass spectrometry and immunoblotting.

### *In vivo* cross-linking and protein complex isolation

Ten million TaC12 cells were harvested and washed with PBS. Subsequently, the cells were incubated with 0.1% (w/v) paraformaldehyde (PFA) in PBS for 8 min at RT to cross-link the proteins. To stop the reaction glycine was added at a final concentration of 125 mM and incubated for 5 min at RT. After this, the cells were washed in PBS. The cell pellets were put on ice and resuspended in ice-cold lysis buffer (20 mM Tris pH 7.5, 140 mM KCL, 1.8 mM MgCl_2_, 0.1% NP-40, 10% glycerol) containing complete protease inhibitor cocktail EDTA free (Roche) and sonicated 3 times for 10 s at 10% power with a Branson Digital Sonifier with 30 s intervals. After centrifugation at 16,000 g for 5 min, the lysate was put on lysis buffer washed Pierce™ Protein A Magnetic Beads (Catalog number: 88845) and Pierce™ Protein G Magnetic Beads (Catalog number: 88847) together with rabbit and rat or mice antibodies, respectively, incubated together prior for 6 h at 4 °C. The lysates were further incubated overnight at 4 °C and subsequently washed three times with RNP lysis buffer. Finally, cross-linking reversal and elution of proteins were performed by incubation with 1 x Lämmli buffer for 20 min at 95 °C and analyzed by mass spectrometry and immunoblotting.

### Mass spectrometry analysis of streptavidin and antibody-based protein pull-downs

Subsequently proteins were shipped on beads for mass spectrometry analysis to the Proteomics and Metabolomics Facility, Center for Biotechnology at the University of Nebraska. Streptavidin bead pull-downs were made up in 50 μL 3x reducing LDS sample buffer containing 15 mM DTT and 2 mM biotin and incubated at 95 °C for 10 min prior to loading all the sample onto a BoltTM 12% Bis-Tris-Plus gel and running briefly into the top of the gel. The gel was fixed and stained with colloidal Coomassie blue G250 stain. Gel containing the proteins was reduced and alkylated, then washed to remove SDS and stain before digestion with trypsin (500 ng) overnight at 37 °C. Peptides were extracted from the gel pieces, dried down, and samples were re-dissolved in 2.5 % acetonitrile, 0.1 % formic acid. 5 μL of each digest was run by nanoLC-MS/MS using a 2 h gradient on a 0.075 mm x 250 mm C18 column feeding into a Q-Exactive HF mass spectrometer. All MS/MS samples were analyzed using Mascot (Matrix Science, London, UK; version 2.6.2). Mascot was set up to search the Bos_taurus_Refseq_002263795.1_ARS-UCD1.2_20190510.fasta (63687 sequences) and cRAP_20150130.fasta (123 sequences) for the three searches, plus one more database for each as described: 1. Old database (OldDB) - Theileria_annulataAnkara_PiroplasmaDB-43_AnnotatedProteins_20190510 database (3796 entries), 2. Uniprot – uniprot-Theileria_annulata_refproteome_UP000001950_20190508 database (3790 entries), 3. New database (NewDB) - Theileria_annulataAnkara_PiroplasmaDB-43_AnnotatedProteins_20191214 database (3572 entries). The searches were done assuming the digestion enzyme trypsin. Mascot was searched with a fragment ion mass tolerance of 0.060 Da and a parent ion tolerance of 10.0 PPM. Deamidated of asparagine and glutamine, oxidation of methionine and carbamidomethyl of cysteine were specified in Mascot as variable modifications. Scaffold (version Scaffold_4.8.9, Proteome Software Inc., Portland, OR) was used to validate MS/MS based peptide and protein identifications. Peptide identifications were accepted if they could be established at greater than 80.0% probability by the Peptide Prophet algorithm (84) with Scaffold delta-mass correction. Protein identifications were accepted if they could be established at greater than 99.0% probability and contained at least 1 identified peptide. Protein probabilities were assigned by the Protein Prophet algorithm (85). Proteins that contained similar peptides and could not be differentiated based on MS/MS analysis alone were grouped to satisfy the principles of parsimony. Proteins sharing significant peptide evidence were grouped into clusters.

For mass spectrometry of proteins that were immunoprecipitated by antibodies as bait, washed protein fractions were loaded on SDS-PAGE gel, fixed in methanol-acetic acid-water (45:1:54) for 20 min and subsequently stained with colloidal Coomassie staining (17% (w/v) ammonium sulphate, 34% methanol, 0.5% acetic acid, 0.1% (w/v) Coomassie blue G-250). Protein bands were excised and stored at 4 °C before analysis. For this, the gel pieces were reduced, alkylated, and digested by trypsin. The digests were analyzed by liquid chromatography LC-MS/MS (PROXEON coupled to a QExactive mass spectrometer, ThermoFisher Scientific, Reinach, Switzerland) with one injection of 5 μL digests. Peptides were trapped on a µPrecolumn C18 PepMap100 (5μm, 100 Å, 300 μm×5mm, ThermoFisher Scientific, Reinach, Switzerland) and separated by backflush on a C18 column (5 μm, 100 Å, 75 μm×15 cm, C18) by applying a 40 min gradient of 5% acetonitrile to 40% in water, 0.1% formic acid, at a flow rate of 350 nl/min. The Full Scan method was set with resolution at 70,000 with an automatic gain control (AGC) target of 1E06 and maximum ion injection time of 50 ms. The data-dependent method for precursor ion fragmentation was applied with the following settings: resolution 17,500, AGC of 1E05, maximum ion time of 110 milliseconds, mass window 2 m/z, collision energy 27, under fill ratio 1%, charge exclusion of unassigned and 1+ ions, and peptide match preferred, respectively. The mass spectrometry data was then searched with MaxQuant (86) version 1.6.14.0 against the following concatenated databases: *Theileria annulata* strain Ankara (PiroplasmaDB, release 43), *Theileria annulata* TaC12 deNovo proteins (manuscript in preparation; no Masking) and uniprot (UniProt Consortium, 2019) *Bos taurus* (release 2021_03), to which common potential contaminants were added. Digestion enzyme was set to trypsin with maximum three missed cleavages, peptide tolerance for first search to 20 ppm and the MS/MS match tolerance to 25 ppm. Carbamidomethylation on cysteine was given as a fixed modification, while methionine oxidation, asparagine, and glutamine deamidation as well as protein N-terminal acetylation were set as variable modifications. Match between runs were allowed between replicates. Identification filtering was controlled by a false discovery rate set at 0.01 at both peptide-spectrum match and protein level. Proteins with one-peptide identification were allowed. Next to MaxQuant’s Label Free Quantification (LFQ) values, protein abundance was also obtained by adding the intensities of the top 3 most intense peptide (Top3) (87), after normalizing the peptide forms by variance stabilization (88). Imputation was performed at the peptide form level for Top3 (iTop3), and at protein level for LFQ (iLFQ). In either case, missing values were replaced by a draw from a Gaussian distribution if there was at most one non-missing value in a group of replicates; this distribution was such that its width was 0.3 x sample standard deviation and centered at the sample distribution mean minus 2.8 or 2.5 x sample standard deviation, for respectively peptide or protein level. Any remaining missing values were imputed by the Maximum Likelihood Estimation (89) method.

### Cell fractionation and isolation of chromatin-bound proteins

Cell fractionation was performed as described above or with Triton X-114 as described previously (54). Briefly 2 ml of Triton X-114 (Fluka, BioChemika, 93422) was resuspended in 98 ml PBS and dissolved at 0 °C and incubated overnight at 30 °C. On the next day the upper aqueous phase was removed and PBS was added to the same volume as removed and again dissolved at 0 °C. This procedure was repeated two more times to obtain 10% condensed Triton X-114. For cell fractionation, a 1mL 1% Triton X-114 solution was made and mixed with pellets of 10 mio TaC12 cells and resuspended by vortexing and sonication 3 x 10 sec at 10% power. After centrifugation at 16,000 g for 5 min, the supernatant was removed and the pellet was put aside. The supernatant was warmed at 37 °C for 1 min (until the solution became cloudy) and then spun at 3000 rpm for 1 min and the upper aqueous phase was removed from the lower detergent phase. Both phases were precipitated by the methanol-chloroform procedure (90) and all fractions were resuspended in 1 x Lämmli buffer. The isolation of chromatin-bound proteins was performed as described previously (57).

### Protein structure predictions

To predict the structure of the identified proteins, IUPred3, flDPnn and AlphaFold2 were used (47, 48, 53, 91). For IUPred3, the analysis type was set to long disorder and medium smoothing. For AlphaFold2, the Multiple Sequence Alignment (MSA) mode utilized was MMseqs2, incorporating UniRef and environmental sample sequence databases (UniRef+Environmental). The number of models was set to 5, with 24 recycles. Convergence of recycles occurred in all models before reaching 24. The relax max iterations were set to 200, and the pairing strategy was set to greedy.

## Funding

This work was founded by the Swiss National Science Foundation (SNF), project number 173972 (P. Olias) and 310030_189127 (S. Rottenberg).

### Author contributions

PO concepted and designed the study. FB, CP, AN, KG, KW and PO acquired and analyzed the data. KW, JR, SR and PO interpreted the data. FB and PO wrote the manuscript with the help of all coauthors.

## Acknowledgements

We would like to thank Brian Shiels for the p104, TashAT2 and Ta9 antibodies. Stephan Grimm is thanked for his technical assistance. Our thanks go to the department for BioMedical Research (DBMR), the Flow Cytometry and Cell Sorting Facility (FCCS) and the Core Facility Proteomics & Mass Spectrometry of the University of Bern, Switzerland. Thanked is the Center for Biotechnology, Proteomics & Metabolomics facility of the University of Nebraska, USA. Jubilee Ajiboye, University of Vermont, is thanked for assistance with alphaFold2.

---

**Figure S1.**
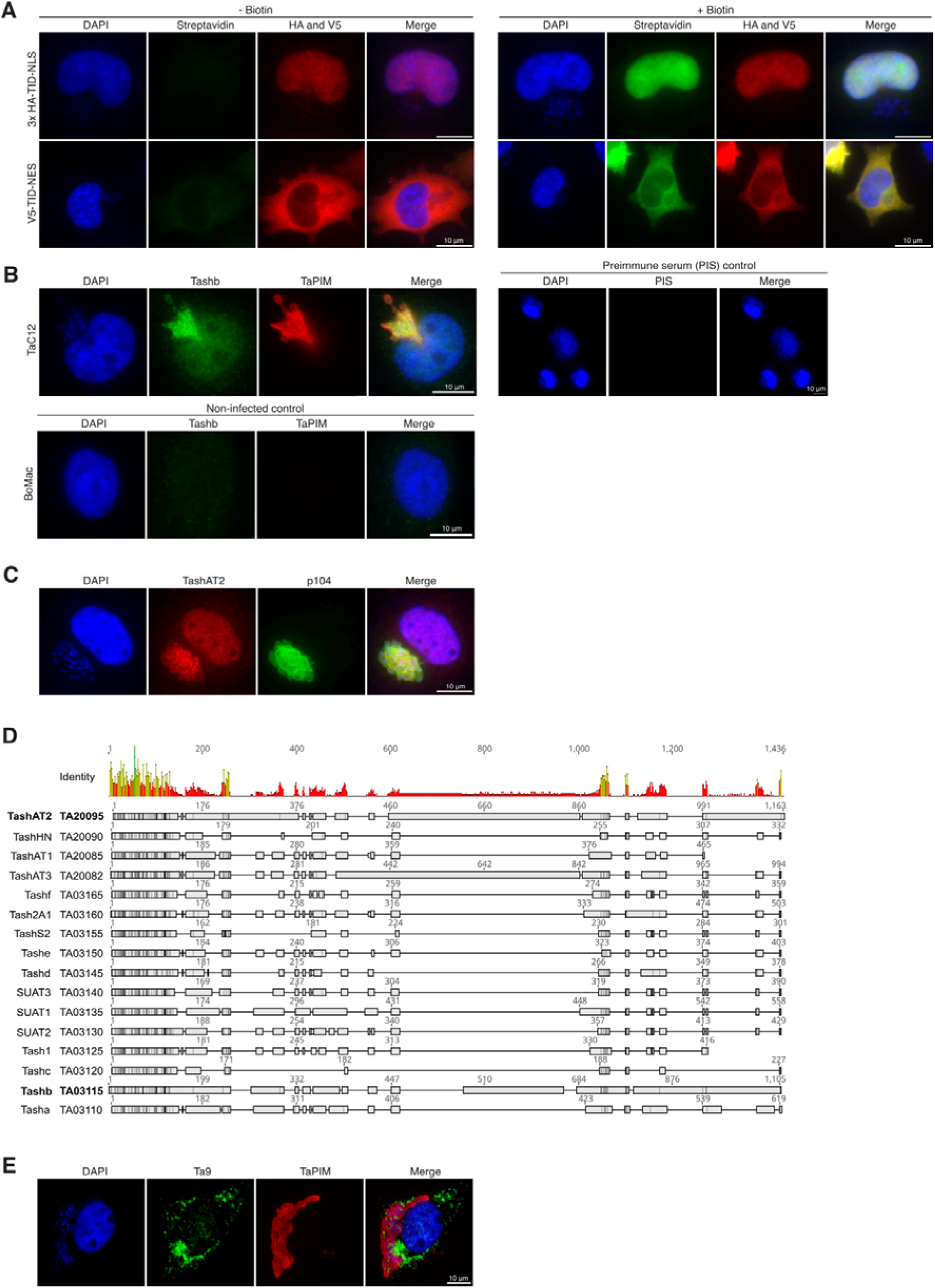
TurboID controls, validation of TashAT2, Tashb and Ta9 protein localization in TaC12 cells, and alignment of Tash and Ta9 locus, related to Figure 1. **(A)** TaC12 cells transduced with HA-TID-NLS and V5-TID-NES constructs, respectively, incubated without (left) and with (right) 150 µM biotin for 30 min prior to fixation. Analysis with α-HA and α-V5 confirms the correct cellular localization of the biotin ligase (red) and enzyme activity (FITC-conjugated streptavidin). Host cell and parasite nuclei are labelled with DAPI. (**B)** TaC12 cells stained with rat α-Tashb (TA03115) and α-TaPIM (schizont membrane). Host cell and parasite nuclei are labelled with DAPI. Right panel shows control stained with pre-immune serum (PIS) of the same rat. Host cell and parasite nuclei are labelled with DAPI. Lower panel shows α-Tashb and α-TaPIM staining of noninfected control cells (BoMac). **(C)** TaC12 cells stained with α-TashAT2 and α-p104 (schizont membrane). Host cell and parasite nuclei are labelled with DAPI. **(D)** Alignment of *T. annulata* TashAT2 and Tashb with the other 14 proteins of the Tash locus. **(E)** TaC12 cells stained with α-Ta9 and α-TaPIM (schizont membrane). Host cell and parasite nuclei are labelled with DAPI.

**Figure S2.**
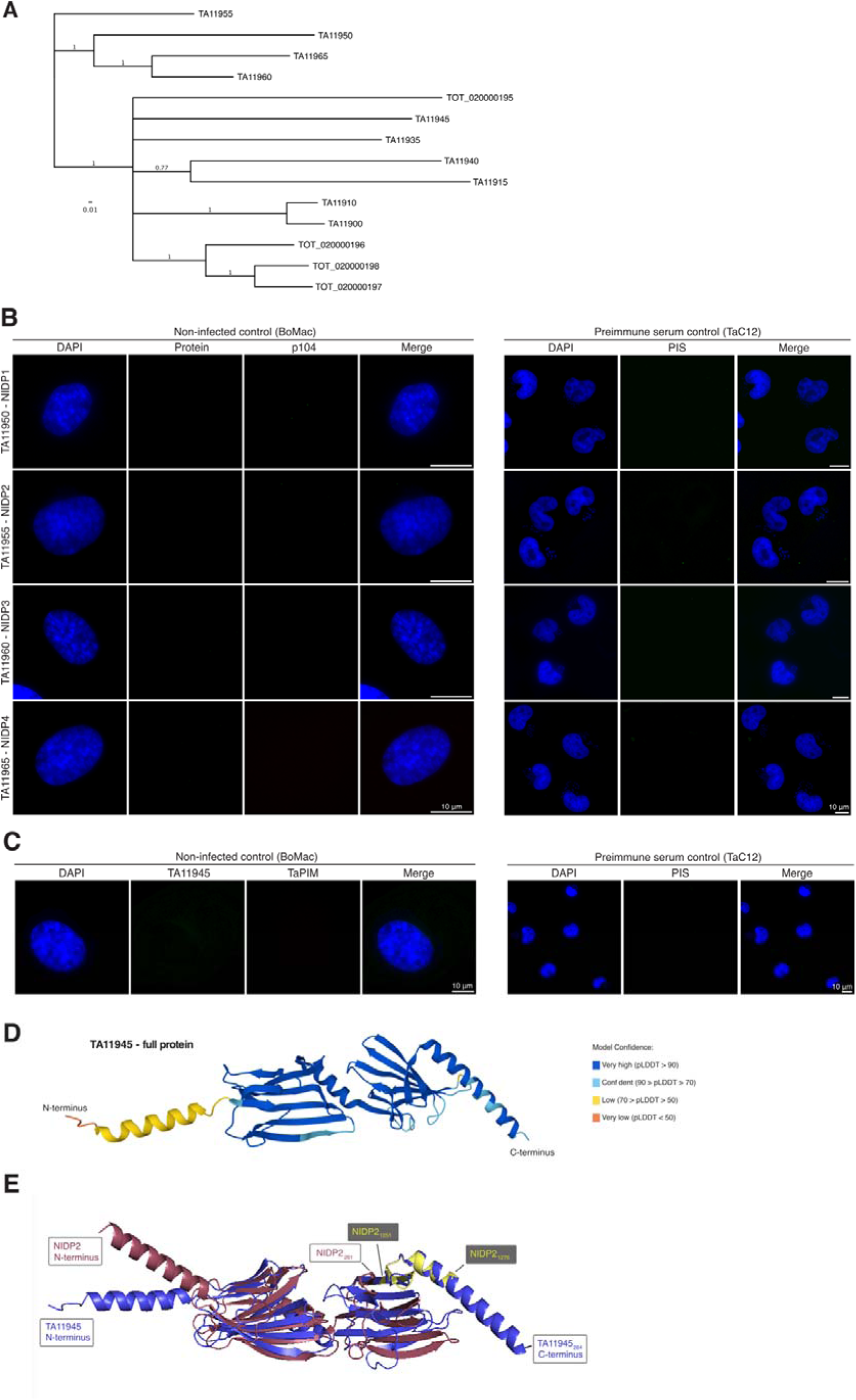
Additional controls and analyses of NIDP1 – 4 and TA11945 proteins, related to Figure 2. (A) Phylogenetic analysis of identified protein family members of *T. annulata* and *T. orientalis*. Note that TA11950, TA11955, TA11960 and TA11965 cluster distinct from *T. orientalis* and other *T. annulata* proteins of this gene cluster, and TA11945 in closest proximity to *T. orientalis* TOT_20000195. (B) As controls, non-infected BoMac cells were stained with α-NIDP1 - 4 and α-p104, and TaC12 cells were stained with corresponding rabbit pre-immune sera (PIS). Nuclei were labelled with DAPI. (C) As controls for TA11945, non-infected BoMac cells were stained with α-TA11945 and α-p104, and TaC12 cells were stained with the corresponding rat pre-immune serum (PIS). Nuclei were labelled with DAPI. (D) Predicted structure of TA11945 by alphaFold2. (E) Overlay of N-terminus and C-terminus of NIDP2 and entire protein structure of TA11945.

**Figure S3.**
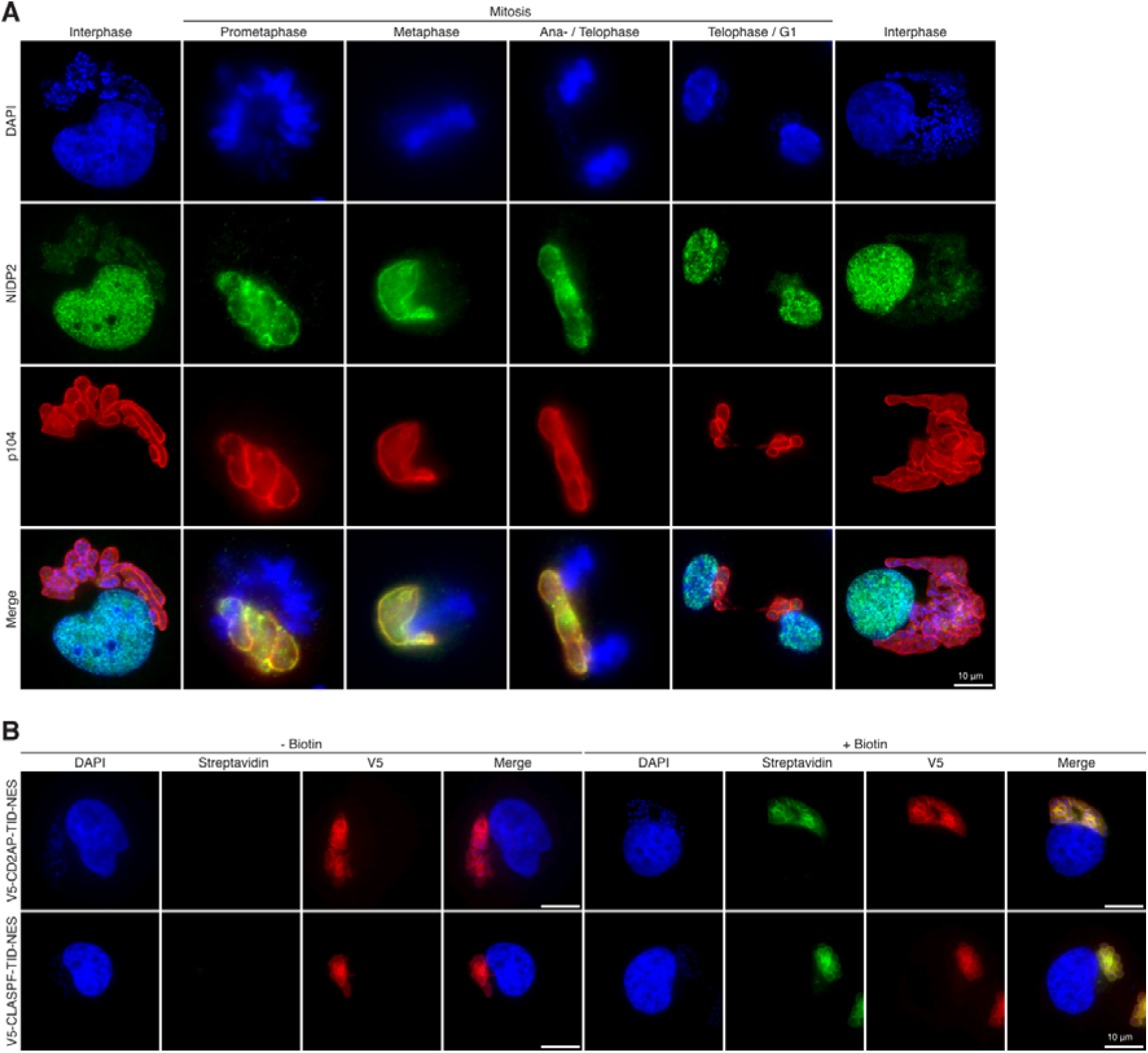
Localization of NIDP2 during interphase and mitosis, and TurboID controls, related to Figure 4. **(A)** TaC12 cells stained with α-NIDP2 in interphase and mitosis. The schizont surface is stained with α-p104, and host and parasite nuclei with DAPI. During prometaphase, metaphase and ana-/telophase NIDP2 colocalizes with p104. In interphase and during telophase/G1 phase NIDP2 localizes in the nucleus of the host cell. **(B)** Transduced cell lines with V5-CD2AP-TurboID-NES and V5-CLASPF-TurboID-NES (CLASPF = CLASP1_1256−1538_) were fixed with PFA and analyzed by immunofluorescence analysis (IFA). V5-tagged-TID was labeled with anti-V5 to confirm the correct localization of the biotin ligase. V5-CD2AP-TurboID-NES and V5-CLASP1_1256−1538_-TurboID-NES cells were incubated with 150 µM biotin for 30 min prior to fixation and analyzed with FITC-conjugated streptavidin. Host cell nucleus and parasite nuclei are labelled with DAPI.

